# Engineering Multiplexed Synthetic Breath Biomarkers as Diagnostic Probes

**DOI:** 10.1101/2024.12.30.630769

**Authors:** Shih-Ting Wang, Melodi Anahtar, Daniel M. Kim, Tahoura S. Samad, Claire M. Zheng, Sahil Patel, Henry Ko, Chayanon Ngambenjawong, Cathy S. Wang, Jesse D. Kirkpatrick, Vardhman Kumar, Heather E. Fleming, Sangeeta N. Bhatia

## Abstract

Breath biopsy is emerging as a rapid and non-invasive diagnostic tool that links exhaled chemical signatures with specific medical conditions. Despite its potential, clinical translation remains limited by the challenge of reliably detecting endogenous, disease-specific biomarkers in breath. Synthetic biomarkers represent an emerging paradigm for precision diagnostics such that they amplify activity-based biochemical signals associated with disease fingerprints. However, their adaptation to breath biopsy has been constrained by the limited availability of orthogonal volatile reporters that are detectable in exhaled breath. Here, we engineer multiplexed breath biomarkers that couple aberrant protease activities to exogenous volatile reporters. We designed novel intramolecular reactions that leverage protease-mediated aminolysis, enabling the sensing of a broad spectrum of proteases, and that each release a unique reporter in breath. This approach was validated in a mouse model of influenza to establish baseline sensitivity and specificity in a controlled inflammatory setting and subsequently applied to diagnose lung cancer using an autochthonous *Alk*-mutant model. We show that combining multiplexed reporter signals with machine learning algorithms enables tumor progression tracking, treatment response monitoring, and detection of relapse after 30 minutes. Our multiplexed breath biopsy platform highlights a promising avenue for rapid, point-of-care diagnostics across diverse disease states.

## Introduction

The COVID-19 pandemic highlighted the critical need for accessible and efficient diagnostics, a demand that now extends to non-communicable diseases. Such tools may offer patients and their physicians the opportunity to personalize decision-making by enabling early detection and longitudinal monitoring of both disease progression and treatment responses. Breath biopsy^1^ has emerged as a promising non-invasive diagnostic approach alongside other minimally-invasive techniques, such as liquid biopsy^2,3^, to facilitate personalized treatment and precision medicine without the need for tissue samples. Volatile organic compounds (VOCs) are exhaled molecules produced endogenously by metabolic processes throughout the body, and have served as breath-based biomarkers for infectious diseases, cancer, asthma, and COPD.^4–8^ Significant advancements in clinical breath biopsy has been made in sampling techniques^9,10^, VOC standardization (e.g., VOC atlas^11^), point-of-care devices^12–16^, such as electronic nose and colorimetric sensors, and machine learning algorithms for analyzing disease-associated VOC patterns.^17^ However, identifying disease-specific VOC signatures remains a major challenge due to heterogeneity in the metabolome profiles, which are highly influenced by an individual’s physiology and environment, resulting in variable breath VOC compositions.^18,19^

Synthetic biomarkers are an emerging paradigm for precision diagnostics, where catalytically-cleavable reporters are designed to probe specific biological activity within diseased microenvironments,^19,20^ and can significantly enhance the signal-to-noise ratios of select analytes. We and others have developed activity-based nanosensors (ABNs), which are diagnostic probes that contain synthetic peptide substrates covalently coupled to exogenous reporters.^21–31^ These probes are modular, such that the readout can be targeted to the activity of specific proteases, or classes of proteases, by changing the substrate sequence, and multiplexed signatures can be generated by incorporating distinguishable reporters. Upon protease cleavage of the ABNs, reporters are liberated from substrates and their relative concentrations in urine have been used to detect cancer, vascular disease, apoptosis, and inflammation.^21–29^ Notably, we previously developed volatile-releasing activity-based nanosensors (vABNs), which are ABNs that generate a breath-based readout through the release of exogenous VOCs. These first-generation vABNs were used to monitor neutrophil elastase activity in bacterial infection and alpha-1 antitrypsin deficiency, by releasing a single volatile fluorinated amine in the exhaled breath.^24^ However, a terminal cleavage reaction was required to release the amine reporters in their native state. This limitation constrained both the target proteases that could be monitored using the vABN platform, and the number of distinguishable volatiles that could be combined into a multiplexed panel. Hence, we were motivated to adopt novel reaction chemistry to create vABNs that could both respond to a broader range of protease classes and release a panel of orthogonal reporters with mass or chemical characteristics that distinguish them from endogenous VOCs, and each other.

In this work, we develop and characterize a multiplexed panel of breath biomarkers through the introduction of a novel mechanism for volatile release from a polymer core, whereby protease cleavage liberates a primary amine that attacks an ester in an intramolecular reaction to generate alcohol reporters (Fig. 1). This self-immolative reaction allows us to selectively probe endopeptidase activity, which hydrolyzes non-terminal amino acids in a peptide substrate. Thus, multiplexed vABNs were generated to sense a broad spectrum of proteases that release volatile amines following exopeptidase cleavage at the terminus and alcohols by endopeptidases. We first validated this approach using a mouse model of viral pneumonia to establish vABN baseline sensitivity and specificity in a controlled inflammatory setting, leveraging well-characterized protease activities associated with viral entry, replication, and host responses.^29,32,33^ We engineered a panel of five vABNs consisting of individual substrates that probe select protease activity and each release a unique amine or alcohol reporter that is detectable in exhaled breath. We showed that multiplexed signals, measured by mass spectrometry and analyzed *via* machine learning algorithms, can accurately differentiate healthy and infected samples. To generalize these findings about chronic disease, we extended this approach to lung cancer detection, treatment response, and relapse. Lung cancer is the leading cause of global cancer mortality, with nearly 70% of patients diagnosed at advanced stages and a five-year survival rate of only 15%.^34^ We re-engineered ABNs that previously contained urinary-based reporters to utilize volatile reporters, and applied them to an autochthonous mouse model of *Alk* mutant non-small-cell-lung cancer (NSCLC).^25,35^ These multiplexed vABNs generated breath-based signals after 30 min of sensor delivery, and distinguished tumor-bearing mice from healthy controls, enabling accurate assessment of treatment responses and post-treatment relapse. Overall, this multiplexed approach for breath biopsy, enabled by an innovative use of chemistry, opens new doors for diagnostics across various pathophysiological conditions.

**Fig. 1:**
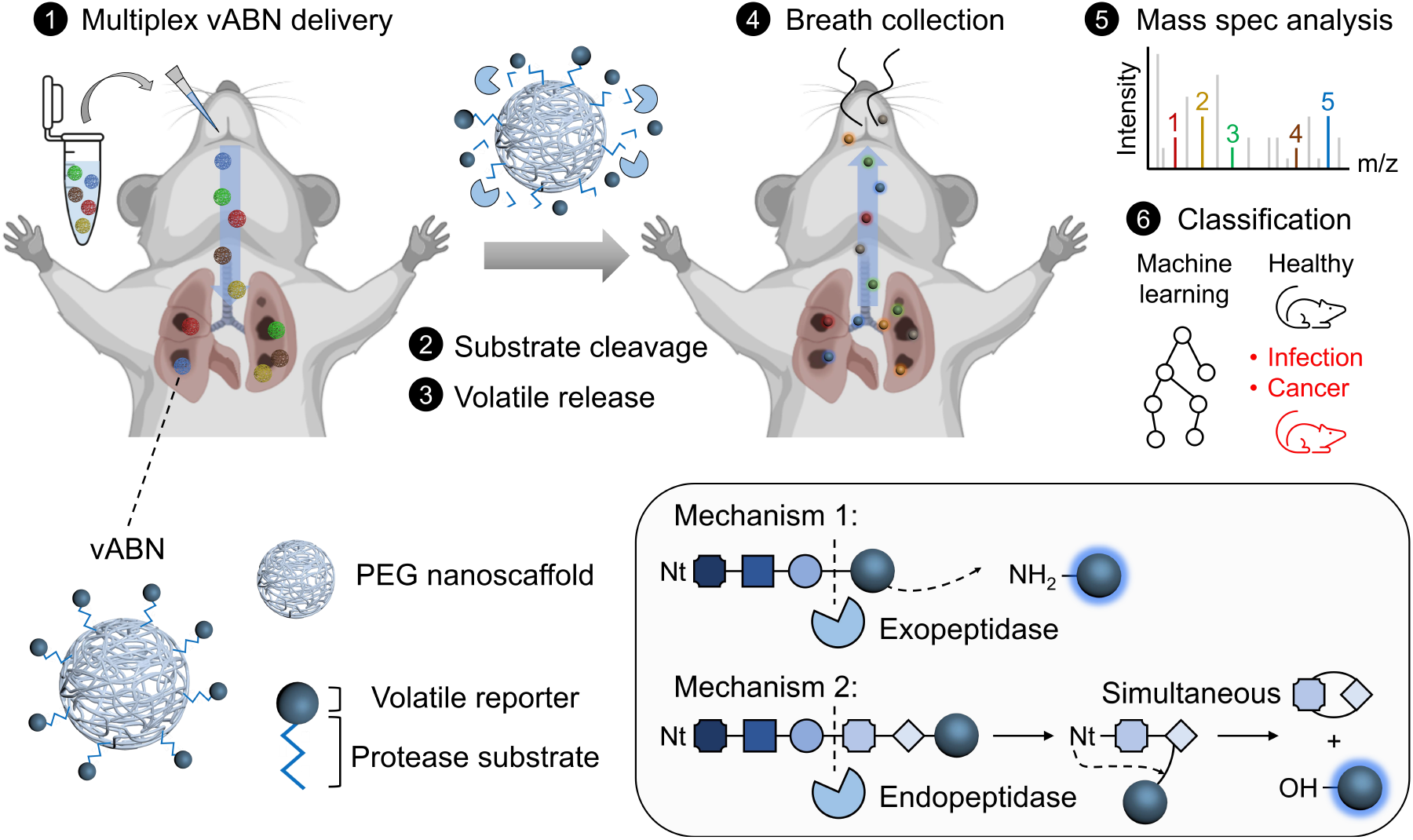
Development of multiplexed breath biomarkers for disease diagnostics. *Top*: multiplexed volatile activity-based nanosensors (vABNs) are delivered locally to mouse lungs, where volatiles are produced, and exhaled breath is assayed by mass spectrometry. Signatures of the detected volatile compounds are used to train classifiers by machine learning algorithms that enable diagnosis of lung infection and lung cancer. *Bottom*: multiplexed vABNs comprised of polyethylene glycol (PEG) nanoscaffolds coupled to volatile reporters through protease-cleavable linkers. Exopeptidase activities are recorded by amine volatiles released through terminal cleavage, and endopeptidase activities by alcohol volatiles *via* controlled aminolysis. Schematic of mice was created in BioRender: Wang, S. (2024) https://BioRender.com/t96g624.

## Results

The ability of a protease to cleave any given substrate is highly dependent on the chemical and spatial arrangements of substrate amino acids relative to the cleavage site (P1/P1’).^36^ For example, we observe significant reduction in the cleavage kinetics of a trypsin-sensing probe, S108^37^, upon substitution of glycine at P1’ to phenylalanine (Supplementary Fig. 1). We sought to generate a volatile-releasing mechanism that preserves cleavage site specificity, in order to detect various types of endopeptidase activities. To this end, we leverage proteolytic cleavage to liberate dipeptide fragments that subsequently undergo intramolecular aminolysis, an established nucleophilic reaction between proximate primary amine and ester groups to generate diketopiperazine and an alcohol.^38,39^ Proline-containing dipeptides are enthalpically favored toward the desired products.^40–42^ Thus, we hypothesized that protease-cleavable probes where proline at P2’ of a substrate was conjugated to an alkyl ester at P3’ would enable selectivity at P1/P1’ following release of an alcohol reporter by aminolysis (Fig. 2a).

**Fig. 2:**
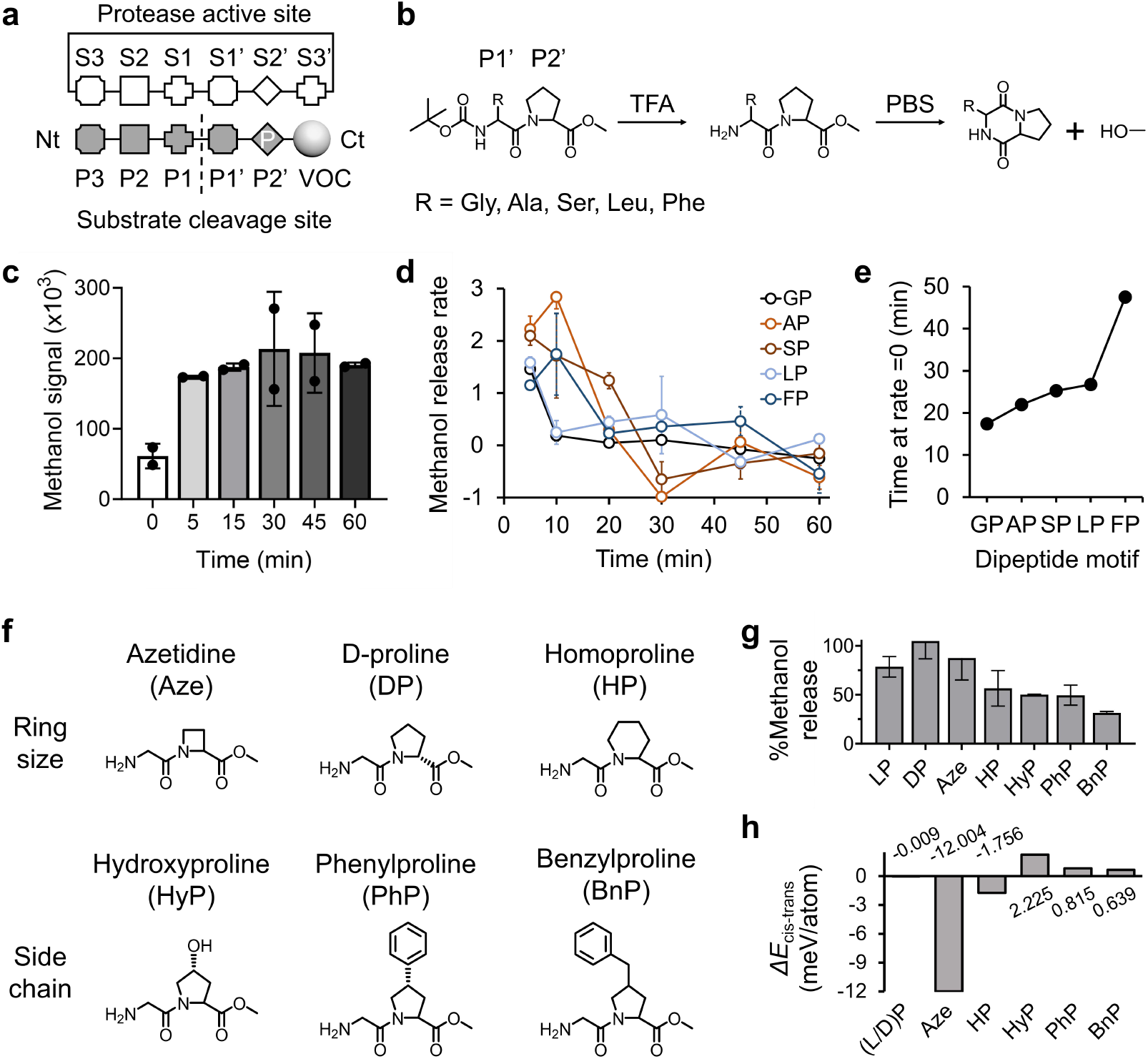
Mechanism of alcohol reporter generation. (a) Schematic showing protease cleavage site and substrate motifs to release alcohol reporters through aminolysis. Nt: N-terminus, Ct: C-terminus, P: proline. (b) Schematic of chemical-induced intramolecular aminolysis of dipeptide methyl ester. Anhydrous trifluoracetic acid (TFA) was used to liberate the *tert*-butyloxycarbonyl (boc) group at the peptide N-terminus. Subsequent reconstitution in phosphate buffered saline (PBS) at 37 °C rapidly proceeds formation of diketopiperazine and methanol. (c) Methanol generated from dipeptide glycine-proline methyl ester (GP-Ome) after chemical cleavage and reconstitution in PBS. (d, e) Kinetics of methanol produced by dipeptides with different P1’ amino acids. (f) Chemical structures of P2’ proline analogs bearing different ring sizes and side chains. (g) Percentage of methanol in the headspace generated by dipeptides with P2’ proline analogs 10 min after reconstitution in PBS. The maximum methanol produced was measured by the addition of tris buffer (50 mM, pH 7.5). (h) DFT analysis of the energy difference between *cis* and *trans* isomers. LP: l-proline, DP: d-proline, Aze: azetidine, HP: homoproline, HyP: hydroxyproline, PhP: 4-phenylproline, BnP: 4-benzylproline.

To investigate the effect of P1’ residue on the kinetics of alcohol release by aminolysis, we synthesized a panel of acid-sensitive *tert*-butyloxycarbonyl (boc)-protected dipeptide methyl esters (Ome), or Xaa-Pro-Ome, where Xaa is glycine (G), alanine (A), serine (S), leucine (L), or phenylalanine (F) in the order of increasing amino acid side chains. These amino acids are also found among the most frequently observed P1’ residues in the MEROPS protease database^43^. The boc group was removed by trifluoroacetic acid, after which methanol generated in the headspace was analyzed by mass spectrometry, following aminolysis of the deprotected dipeptides in phosphate buffered saline at pH 7 at 37 °C (Fig. 2b and Supplementary Fig. 2). As shown in Fig. 2c, deprotected GP-Ome exhibits rapid generation of methanol and plateaus within 20 min, suggesting equilibrium in the gas phase. The relative ranking of methanol release rates is in the order of G>A>S>L>F, such that the rate decreases with increasing size of the side chains (i.e., amount of time needed to reach rate=0; Figs. 2d and e). Regardless of the size of P1’ residue, reactions were completed within 1 hour. Notably, non-specific hydrolysis at the ester group was absent in common buffers at physiological pH of 5.5, 7, and 9, except when using a high concentration (50 mM) of amine-containing tris buffer (Supplementary Fig. 3). These observations suggest that using an aminolysis reaction to achieve volatile release is compatible with use in a diagnostic test.

We further investigated the effects of P2’ by substituting proline with its analogs (Fig. 2f), which are frequently used to modulate protease substrate specificity. Previous studies have shown that substituents on the pyrrolidine ring of proline residues have significant structural consequences.^40,42,44^ Our results align with these findings, showing rapid methanol release, particularly from dipeptides containing 4- or 5-member rings at P2’, but not from those with side chains on the pyrrolidine rings (Fig. 2g). We also validated the energy difference between *cis-trans* conformers of each dipeptide using density functional theory (DFT) calculations, which suggested that efficient aminolysis is facilitated by a preference for *cis* conformation (Fig. 2h).

We previously reported a volatile sensing activity-based nanosensor (vABN)^24^ composed of an eight-arm polyethylene glycol (PEG) nanoscaffold attached to a protease substrate and exogenous reporter (Fig. 3a). To examine alcohol release following selective protease cleavage, we modified an established trypsin-cleavable substrate from our database, S108^37^, with proline and methyl ester. The resulting modification, designated S108-Ome (Ac-GAANLTRGP-Ome), should yield a substrate that can be canonically cleaved by human Trypsin-3 (PRSS3) after the arginine residue, and result in methanol release by aminolysis. The molecular weights of any intact peptide and cleaved fragments present in the liquid phase were monitored by MALDI-TOF mass spectrometry (Fig. 3b), while volatile methanol generated by aminolysis of fragment GP-Ome was detected in the headspace (Fig. 3c). To demonstrate that alcohol release is dependent on cleavage site specificity, we incubated S108-Ome with kallikrein-14 (Fig. 3c, bottom), but in this reaction, no methanol was detected (Fig. 3d) due to this enzyme utilizing a different cleavage site between P1 asparagine and P1’ threonine (Fig. 3b). The selectivity towards target proteases could be further enhanced by substituting l-proline with d-proline at P2’, which prevented non-specific release of alcohols by prolyl peptidases, like fibroblast activation protein (FAP), which is involved in inflammation and cancer^45^ (Figs. 3e and f). To prevent off-target cleavage, the d-proline at P2’ was used in all subsequent vABNs in this work. Further, methanol release was well correlated with PRSS3 activity, which is known to be active at neutral and basic pH but not in acidic conditions (Fig. 3g).

**Fig. 3:**
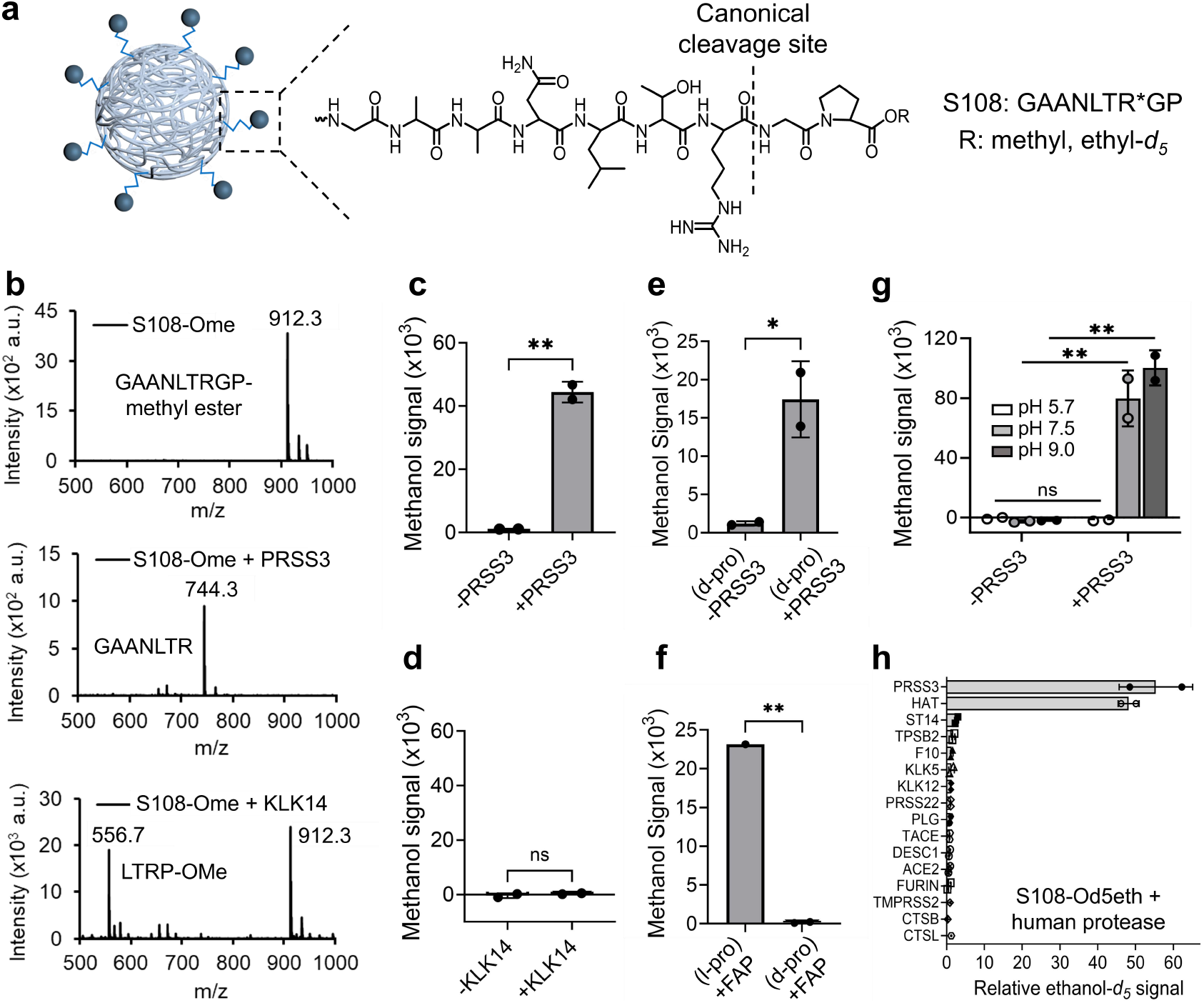
Volatile detection mediated by protease cleavage. (a) Schematic of a vABN where trypsin-sensing substrate S108 is attached to volatile reporters. The dotted line and * indicate the canonical cleavage site that initiates alcohol release by aminolysis. R indicates alkyl group. (b) MALDI-TOF spectra of S108 methyl ester (S108-Ome) before (top) and after incubation with human trypsin 3 (PRSS3, middle) or kallikrein-related peptidase 14 (KLK14, bottom) for 1 hr at 37 °C. (c, d) Methanol signal produced by S108-Ome with PRSS3 or KLK14. (e) Methanol signal generated by S108(d-pro)-Ome with PRSS3, where P2’ is a d-proline. (f) S108(l-pro)-Ome and S108(d-pro)-Ome were used to assess non-specific cleavage by fibroblast activation protein (FAP). (g) S108-Ome was used to assess PRSS3 activity modulated by solution pH. (h) Ethanol-*d*_*5*_ signal generated by screening S108 ethyl-*d*_*5*_ ester (S108-Od5eth) against 16 recombinant human proteases associated with influenza A viral infection. Significance was calculated by two-tailed unpaired t tests (c-f) and two-way ANOVA with Bonferroni corrections for multiple comparisons (g); *P < 0.05, **P < 0.01.

Trypsins, such as PRSS3, are known to be involved in the host entry and replication of influenza A virus.^32,33^ To demonstrate that S108 activity is specific to trypsin-like proteases, we screened S108 ethyl-*d*_*5*_ ester, which can release biocompatible ethanol-*d*_*5*_, against PRSS3, the closely-related human airway trypsin-like protease (HAT), and 14 other non-trypsin recombinant human proteases, and only observed meaningful alcohol release in response to trypsin activity (Fig. 3h). To test the ability of S108 vABNs to detect infection-driven trypsin activity we generated an Influenza A (A/Puerto Rico/8/34, PR8) mouse model according to a method used in our recent studies^29^ (Fig. 4a). We collected longitudinal samples of bronchoalveolar lavage fluids (BALF), which act as a proxy for the lung airspace, from PR8-infected mice. Over the course of infection, we detected elevated trypsin levels by enzyme-linked immunosorbent assays (ELISAs) and increasing trypsin activity using S108-Ome in BALF (Figs. 4b and c). We then sought to test whether these BALF signals correlated to breath by intratracheally delivering S108 vABNs to healthy and PR8-infected mice on day 6 post-infection. Breath collected 10 min post-vABN delivery showed increased levels of ethanol-*d*_*5*_ in infected mice *versus* healthy controls (Fig. 4d), suggesting that the S108 vABN could be used to non-invasively detect trypsin activity that is associated with viral infection.

**Fig. 4:**
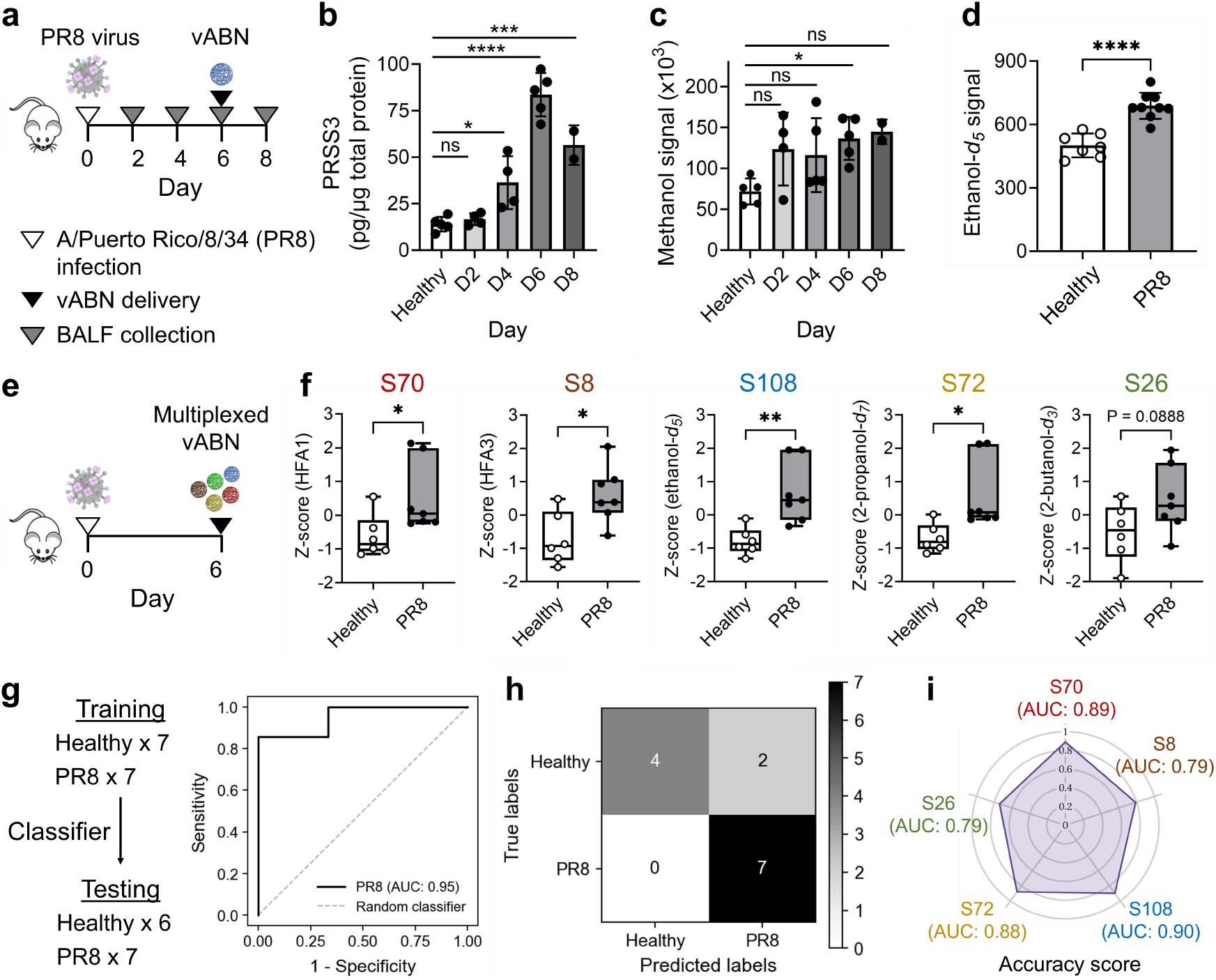
Development of multiplexed vABN *in vivo*. (a) Schematic of the influenza mouse model, where a single plex vABN was delivered 6 days post-infection. (b) Mouse trypsin (PRSS3) protein levels were quantified by ELISA of BALF from healthy and PR8-infected mice collected at 2-, 4-, 6-, and 8-days post-infection. (c) Trypsin-sensing vABN, S108-Ome, monitored progression of lung infection by detecting methanol released into BALF. (d) vABN-derived ethanol-*d*_*5*_ levels were measured in exhaled breath of healthy and D6 PR8-infected mice at 10 min after S108 vABN delivery. (e) Schematic of approach. Multiplexed vABNs, S70 (HFA1), S8 (HFA3), S108 (ethanol-*d*_5_), S72 (2-propanol-*d*_7_), and S26 (2-butanol-*d*_3_) were delivered to healthy and PR8 mice 6 days post-infection. (f) Standardized volatile signals by z-scoring individual vABNs in the 5-plex panel, based on measurements in exhaled breath. (g) A random forest classifier was applied to independent training and testing groups of healthy and PR8 mice (n=6 or 7 mice in each group). Performance was represented with an ROC curve and estimates of out-of-bag error were used for cross-validation. (h) Distinguishing power of the trained classifier was visualized with a confusion matrix to differentiate healthy from PR8 mice. (i) A radar chart showing AUC values of individual vABNs in the multiplexed panel. Significance was calculated by one-way ANOVA with Bonferroni corrections for multiple comparisons (b, c) and two-tailed unpaired t tests (d, f); *P < 0.05, **P < 0.01, ***P < 0.001, ****P < 0.0001.

Having established a novel volatile release mechanism that can generate breath-based disease signal, we next sought to test the feasibility of multiplexing vABNs in breath. A panel of five vABNs was constructed, each containing a different peptide substrate-reporter pair. Two of the vABNs were synthesized with the reporters pentafluoropropylamine (HFA1) and perfluoroamine (HFA3), which have terminal amines that are compatible with the vABN chemistry previously demonstrated to sense exopeptidase-like activity^24^. The other three vABNs contained alcohol reporters, including ethanol-*d*_*5*_, 2-propanol-*d*_*7*_, and 2-butanol-*d*_*3*_ for endopeptidases. Four substrate sequences (S8, S108, S72 and S26, Supplementary Table 1) was selected from our database^37^ and were modified to increase susceptibility to influenza-associated proteolytic host responses. This panel also included S70, also known as BV01, which is susceptible to cleavage by granzyme B activity and that we recently identified as a highly sensitive probe for distinguishing host responses in viral *versus* bacterial pneumonia.^29^ The 5-plex vABN cocktail was delivered directly into the lungs of PR8-infected mice, and breath was collected 10 mins later. Multiplexed signals were standardized *via* z-score and subjected to machine learning analysis using a random forest classifier (Figs. 4e and f). Diagnostic accuracy was measured by the receiver operating characteristic (ROC) curve, which distinguished infected mice from healthy controls with an area under the curve (AUC) of 0.95 and an out-of-bag error rate of 2.14%, suggesting high accuracy without significant overfitting (Fig. 4g). Performance of the trained classifier was also visualized with a confusion matrix (Fig. 4h). The AUC values from individual vABNs suggested that multiplexing enhanced overall accuracy, as the combined diagnostic power outperformed individual sensors alone (Fig. 4i).

Having established that 5 distinct VOC reporters can be analyzed collectively to accurately detect active influenza infection, we sought to apply this multiplexing approach for the detection of lung cancer, the global leading cause of cancer mortality.^34,46,47^ To address this clinical need for diagnostic and monitoring tools, our group previously created a multiplexed panel of 14 activity-based nanosensors (ABNs) capable of noninvasive monitoring of lung cancer *via* a urinary readout.^25,35^ These 14 ABNs were designed by *in vitro* screening of a substrate panel against the activity of enzymes from classes associated with lung cancer, including serine and metalloproteases. Critically, this panel included peptide substrates recognized *in vitro* by a range of proteases that released mass-barcoded urinary reporters following substrate cleavage.^25,26^ In an *Alk* mutant (Eml4-Alk) model, this 14-plex of urinary ABNs could differentiate healthy and tumor-bearing mice, and monitor response to treatment by alectinib, a first-line ALK tyrosine kinase inhibitor (TKI) for advanced *Alk*^+^ NSCLCs.^48^

To adapt this platform as a breath-based diagnostic, we re-engineered a subset of the urinary ABNs (PP01, 9, 10, 12, and 13) into a new 5-plex vABN panel. First, to validate that the updated substrates can release volatile reporters following proteolytic cleavage, we modified each sequence with methyl ester, incubated them in BALF collected from healthy and Eml4-Alk mice at 7 weeks post tumor initiation, and measured methanol concentration in the reaction headspace. We detected differential signals between healthy and tumor-bearing mice, particularly for sensors PP09 and PP10, which were designed to detect activity of MMPs and serine proteases, respectively (Supplementary Fig. 4a and Supplementary Table 1). After confirming that these substrates were capable of volatile release, we synthesized them into multiplexed vABNs. These five vABNs use the same volatile reporters as the PR8 5-plex, but contain entirely different substrate sequences, thus constituting novel substrate-reporter pairs (Supplementary Table 1). Micro-computed tomography (microCT) scans were obtained prior to all breath collections. Multiplexed vABNs were delivered to the lungs of healthy and Eml4-Alk mice at 4-, 5-, or 6-weeks after tumor initiation (Fig. 5a). Breath was collected 10 and 30 mins after vABN delivery and volatile signal amplitudes were integrated over 30 min by area under the curve analysis (AUC). The multiplexed signals generated within 30 min post-vABN delivery were standardized and subjected to machine learning analysis using the gradient boosting algorithms^49^. Diagnostic accuracies were measured by ROC curves, which distinguished Eml4-Alk mice from healthy controls with combined AUCs of 0.64 at 4 weeks, 0.79 at 5 weeks, and 0.93 at 6 weeks (Fig. 5b). Five-fold cross validations were performed to evaluate the classifiers, of which mean AUCs, F1 and precision scores were reported in Supplementary Table 2. The AUC values of individual vABNs suggested that multiplexing enhanced the overall accuracy (Fig. 5c) of the diagnostic.

**Fig. 5:**
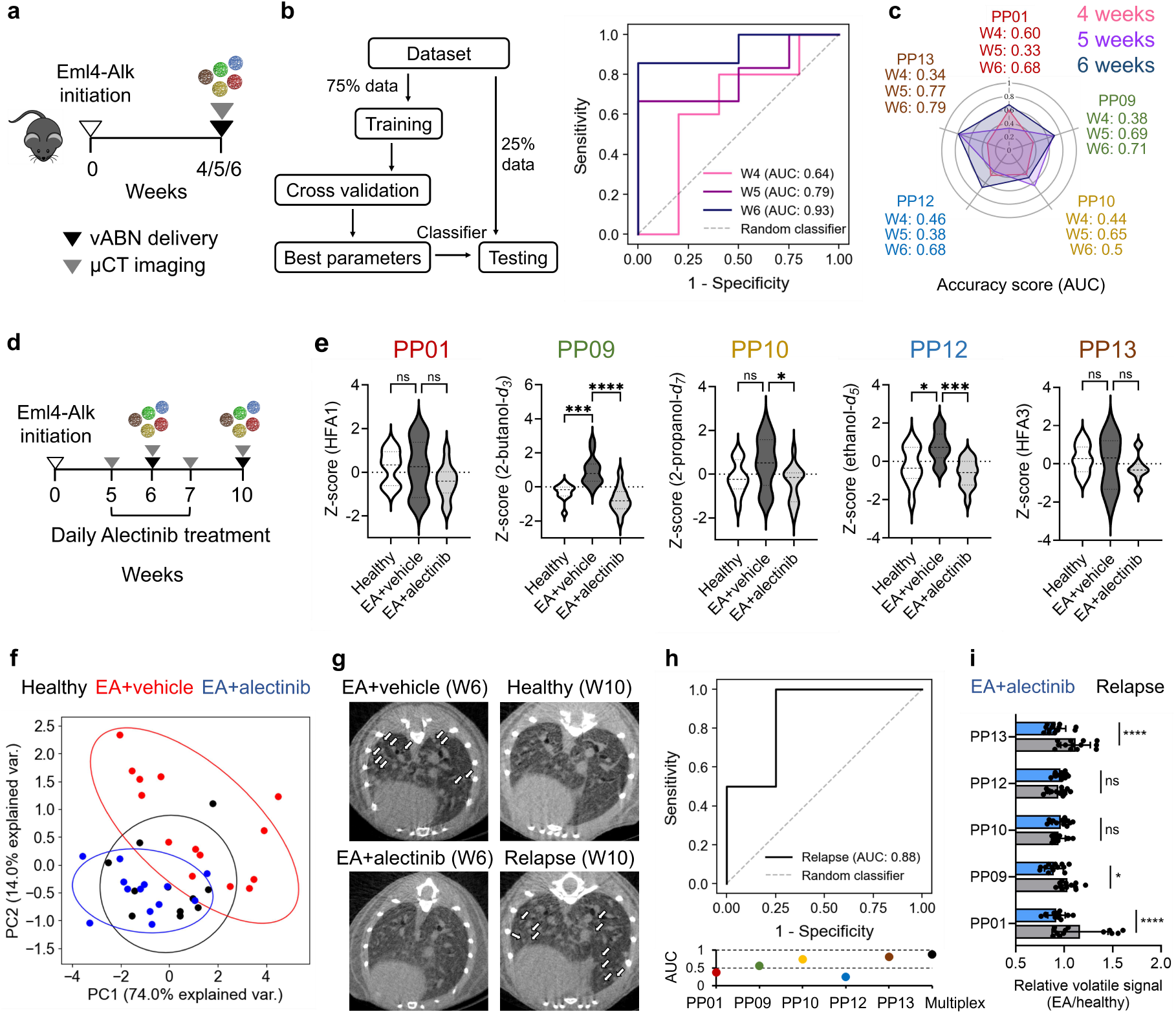
Multiplexed vABNs for lung cancer detection. (a) Schematic of the Eml4-Alk (EA) model, where multiplexed vABNs were delivered at 4-, 5- or 6-weeks post tumor initiation. (b) ROC curve representing the performance of a gradient boosting classifier in differentiating EA mice from healthy controls. The data was concatenated from two independent standardized cohort signals (healthy: n = 15, 16, 14 and EA: n = 25, 24, 21 for 4, 5, and 6 weeks, respectively). (c) A radar chart showing AUC values of individual vABNs in the multiplexed panel at different weeks. (d) Schematic of approach. Multiplexed vABNs were delivered to mice one week after alectinib treatment and three weeks after discontinuing the treatment. (e) Violin plot summarizing the volatile signals of individual vABNs in the 5-plex panel among the three states of interest at week 6 post tumor initiation (healthy: n = 11, EA+vehicle: n = 14, EA+alectinib; n = 13). (f) PCA analysis was performed on standardized vABN signals from healthy controls (black) and EA mice treated with either vehicle (red) or alectinib (blue). (g) Representative microCT scans of representative lung tissues from mice after one week with or without alectinib exposure (W6), or three weeks after discontinuation of treatment (W10). White arrows point to tumors. (h) ROC curve (top) and AUC values (bottom) show performance of a gradient boosting classifier in differentiating relapsed EA mice (n = 13) from healthy controls (n = 11) at W10. (i) Individual vABN signals at 6 weeks (with alectinib treatment; blue bars) and at week 10 (three weeks after discontinuing the drug; grey bars). The raw signals were normalized to the mean signals of healthy controls in the same cohort at the same week. Significance was calculated by one-way (e) or two-way (i) ANOVAs with Bonferroni corrections for multiple comparisons; *P < 0.05, **P < 0.01, ***P < 0.001, **** P < 0.0001.

In addition to monitoring for the growth of lung tumors, multiplexed vABNs detected altered protease activity patterns in mice following treatment with alectinib (Figs. 5d and e). Principle component analysis (PCA) revealed that protease activity in vehicle-treated Eml4-Alk mice diverged from healthy controls, whereas that of alectinib-treated mice were more similar to healthy controls (Fig. 5f). We also explored the MMP specificity of PP09 by incubating methanol probe PP09-Ome with recombinant MMP9, MMP12, and MMP13, which are elevated in the transcriptomic data of Eml4-Alk mice^25^. As shown in Supplementary Figs. 4b-g, elevated methanol signal was observed specifically for MMP9, which agreed with the elevated protein levels in BALF quantified by ELISAs.

The effective regression of tumors by alectinib treatment was observed by microCT scans (Fig. 5g top). In clinical trials, high initial response rates to tyrosine kinase inhibitors have been observed for this patient population; however, many patients acquire resistance to TKIs and ultimately relapse on therapy within 1 to 2 years.^50,51^ We therefore sought to assess our approach in detecting tumor regrowth. To this end, alectinib treatment was terminated after two weeks, and mice were monitored by microCT and vABN. As indicated by microCT lung scans, mice treated with alectinib that previously bore tumors exhibit tumor regrowth (an experimental proxy for clinical relapse) three weeks after discontinuation of the drug (Fig. 5g bottom). Similar to the approach above, multiplexed vABNs were delivered to healthy and relapsed mice, and the standardized volatile signatures of each group were used to train classifiers. As shown in Fig. 5h, our results showed high accuracy of classification, offering a feasible method to monitor cancer relapse. Furthermore, multiple vABNs (PP01, PP09, PP13) exhibited marked signal changes following relapse compared to those observed during alectinib treatment, indicative of altered pulmonary protease activity in relapsing tumors (Fig. 5i).

## Discussion

In this study, we design and credential a novel chemistry-based approach to multiplex breath-based synthetic biomarkers by coupling disease-associated protease activity with distinct exogenous volatile reporters. This multiplexed volatile activity-based nanosensor (vABN) platform, when combined with analysis by machine learning algorithms, accurately classified disease states in preclinical models of viral infection and lung cancer using rapid breath tests. Importantly, we extended this approach to monitor both treatment response and relapse in a mouse model of *Alk*-mutant lung cancer, highlighting the modular nature of vABNs, and their suitability for adaptation to various pathophysiological processes that currently require monitoring by invasive procedures.

The vABN developed here is comprised of a nanoscaffold attached to protease-sensing substrate and exogenous reporter. To achieve multiplexing in breath, the core innovation lies in the ability to detect dysregulated protease activities of broad catalytic types. Although the incorporation of current self-immolative linkers^52^ can release volatile amines and alcohols through proteolytic cleavage at the terminus, effective mechanisms for sensing endopeptidase activity, which accounts for >50% of known 589 human proteases^43,53,54^ have remained out of reach. To fill this gap, we developed a novel reaction to release volatile reporters upon selective endopeptidase cleavage, including several disease-associated metallo- and serine proteases. The mechanism worked by coupling protease scission with conformational-constrained motifs to facilitate rapid head-to-tail cyclization of proline-containing dipeptide fragments *via* aminolysis, along with alcohol production. Although intramolecular aminolysis^38–42^ is well-established, to our understanding, it is the first time that this reaction has been used as a volatile release mechanism for breath readouts. Dipeptides containing the most common P1’ amino acids, including glycine, alanine, serine, and leucine, complete reactions within 30 min, enabling rapid breath tests. We also identified proline analogs, including d-proline, azetidine, and homoproline, that promoted the *cis*-configurations required for intramolecular aminolysis, thereby expanding the scope of detectable protease activities *in vivo*.

The first test of this platform was to apply it to an influenza mouse model by designing a 5-plex panel that was capable of detecting protease activities associated with viral infection. Each vABN was attached to either an amine- or alcohol-based reporter that was distinguishable by mass and did not exist in the breath. This multiplexed panel included several novel probes, highlighting S108 that was demonstrated to be cleaved specifically by trypsin in this work, and S70 (or BV01), which was re-engineered from our previously identified urinary ABN for host granzyme B activity that distinguished viral *versus* bacterial pneumonia.^29^ In this context, these two vABNs were individually able to classify infected mice from healthy controls with AUC values of ∼0.90, and performance improved further when multiplexed (AUC of 0.95). We then extended this platform to an *Alk*-mutant lung cancer model by generating an unique set of 5-plex vABN panel adapted from previous urinary ABNs that differentiated tumor bearing mice from healthy controls^25,26^. This re-design of ABNs with volatile reporters would not have been possible using the first-generation vABN we published previously, as it only detects exopeptidase activities *via* terminal cleavage and the most predictive host response was driven largely by endopeptidase activity. Analysis of multiplexed signals by machine learning algorithms provided accurate assessments of tumor progression and treatment response 30 min post administration with enhanced classification performances compared to single vABNs. These results demonstrate modularity of the protease-sensing vABN platform to diagnose complex disease with nuanced signature *via* breath.

Among the five vABNs, volatile signals from PP09, PP10, and PP12 vABNs exhibited an overall increase in the vehicle-treated mice and regression with alectinib treatment. Notably, the responses of PP09 and PP10 agreed with previous urinary diagnostics of lung cancers^25,26^ but not PP12. We hypothesize that PP12, now with a cleavage site at arginine for serine proteases, may be cleaved by distinct proteases (e.g., MMP) in the urinary panel; however, further study would be required to explore this. Overall, the MMP-specific PP09 was the most sensitive vABN for monitoring tumor progression (Figure 5c) and treatment-related changes (Figure 5e). As an endopeptidase, our novel volatile-releasing chemistry allowed for identifying cleavage sensitivity and site specificity to PP09 by subclasses of tumor-associated MMPs. These findings demonstrate the modularity and complementary nature of ABN and vABN platforms when used for multimodal readout in urine and breath. The multiplexed vABN platform was further applied to monitoring for recurrence of *ALK*^*+*^ NSCLCs, as a crude proxy of relapse driven by clinical resistance to first-line ALK inhibitors.^50,51^ Clinical resistance to TKIs can occur through a variety of mechanisms. As a proof-of-concept to model this outcome, we discontinued alectinib treatment of mice, and functional relapse of the animals was scored based on tumor regrowth, and which was universally observed within three weeks after drug withdrawal. Multiplexed vABNs (PP01, PP09, PP13) detected elevated protease activities in mice that relapse with tumor growth following the withdrawal of alectinib administration, supporting the clinical potential of this approach for monitoring treatment efficacy. Notably, PP13, engineered for cleavage by serine proteases, was not only the key feature driving classification between healthy and relapse mice (Figure 5h), but also exhibited marked signal changes between relapse and treatment (Figure 5i). These observations suggest novel protease activity that may distinguish primary tumor growth from relapse and warrant further investigation.

Modular diagnostic systems offer the opportunity for future optimization across many dimensions: 1) nanosensor formulation and molecular targets, 2) route of administration and reporter collection, and 3) reporter chemistry and analytics for readout. In this work, we describe features of one particular system, employing a peptide-based formulation to monitor a panel of proteases, a pulmonary delivery method, a breath-based reporter collection, and a readout using PTR-MS (a type of soft ionization mass spectrometry). Other published reports have highlighted different formulations, delivery, collection methods, and readouts. In considering sensor formulations, many enzymatic hydrolysis can potentially be leveraged.^20,22^ For example, aside from proteases, glycosidases have emerged as another promising class of targets for ABNs, with corresponding formulations based on sugar molecules^55–58^. These sugar-based ABNs can release ethanol-*d*_5_ as a readily translatable VOCs and is currently being evaluated in a phase 2 clinical trial. In our work, novel aminolysis chemistry leverages peptide-based vABNs to also release alcohol VOCs. In comparing enzyme classes, proteases tend to have equivalent or higher catalytic efficiency (k_cat_/k_m_) than glycosidases. Further, there are significantly more known proteases^59^ than glycosidases^60^ in the human genome, and the protease biology of disease is currently more advanced than glycosidase biology although this field is rapidly evolving.^59,60^ These guide our ABN-based tests towards leveraging proteases for disease applications. In terms of delivery, systemic administration, such as intravenous injection can query a broad range of tissues but is limited by off-target activity in the blood and other organs. Intratracheal delivery achieves enrichment for pulmonary signals but is currently limited to detection of pulmonary diseases. The addition of targeting moieties such as nanobodies and peptides can further enhance both sensitivity and specificity of activity-based sensors.^23,31^ Finally, in considering the selection of VOCs, we note that the current 5-plex is limited by exogenous reporters that can be distinguished by mass using PTR-MS. This method offers near real-time analysis and low limit of detection, but a reduced chemical resolution compared to a hard ionization technique such as gas chromatography mass spectrometry (GC-MS). Future efforts could focus on expanding the chemical space of reporters by employing GC-MS coupled with auto-sampling platforms.

Aside from tuning exogenous vABN components, a potential future strategy for increasing diagnostic accuracy includes analyzing the synthetic vABN signals together with endogenous VOCs with machine learning. In addition, multimodal diagnostics that combine ctDNA, protein biomarkers, and/or circulating tumor cells may enhance diagnostic accuracy over any one modality alone. Finally, as breath analysis devices continue to be miniaturized and their resolution improved, such as for 3D compact mass spectrometers or portable Raman spectrometers, integrating this vABN technology with such devices could facilitate its implementation in point-of-care diagnostics, ultimately contributing to earlier detection, better patient stratification, and improved outcomes in protease-mediated diseases.

## Methods

### Peptide synthesis

*tert*-Butyloxycarbonyl (boc) protected dipeptide alkyl esters and protease substrate alkyl esters were synthesized by CPC Scientific or the Biopolymers and Proteomics Core at the Swanson Biotechnology Center at the Koch Institute of MIT. Briefly, peptides were synthesized by standard solid phase synthesis. To couple desired alcohol reporters to the peptides, Steglich esterification^61^ was first used to prepare boc-Yaa-alkyl ester, where Yaa is proline or its derivatives. Boc was removed by anhydrous trifluoroacetic acid (TFA) followed by coupling to the remaining peptide fragment. All protecting groups were removed in final TFA treatment. Peptide products were purified by high performance liquid chromatography (HPLC) and analyzed by matrix-assisted laser desorption/ionization time-of-flight (MALDI-TOF) mass spectrometry.

### Density Functional Theory (DFT) calculations

The energy of dipeptide conformers was calculated using ORCA (v5.0). The initial geometry was generated by ChemDraw (20.1.1, Revvity Signals) and optimized using the built-in Universal Force Field in Avogadro (v1.2.0). DFT geometry optimization was then performed with the B3LYP functional. We applied the def2-TZVP basis set, which provides triple-zeta quality and polarization functions for greater accuracy in representing electron distributions. The RIJCOSX approximation, supported by the def2/J auxiliary basis set, was used to speed up the calculation of the Coulomb and exchange integrals through a resolution of the identity (RI) approach. The KDIIS algorithm was applied to enable efficient convergence of the self-consistent field (SCF) procedure. Finally, Grimme’s D3BJ dispersion correction was included to account for *van der* Waals interactions, which are often underestimated in DFT calculations.

### *In vitro* headspace analysis

Aminolysis reaction in Fig. 2 was performed as follows. Boc-protected dipeptide methyl esters were dissolved in dichloromethane (DCM) and aliquoted to ∼10 µmol in 1.7 mL tubes. After evaporating DCM, anhydrous 30% trifluoracetic acid (TFA) in DCM was incubated with the peptides for ∼3 hrs at room temperature to deprotect the boc group. After evaporation of TFA and DCM, 100 µL of phosphate buffer (50 mM) was added to the deprotected dipeptides and adjusted to pH∼7 with 10 M sodium hydroxide. In Figure 3, protease cleavage mediated aminolysis was performed by incubating peptide alkyl esters (100 µM) with target proteases (100 nM, R&D Systems or Enzo Life Sciences) in a total volume of 100 µL for 1 hr at 37 °C. The released alcohols in the headspace were analyzed by proton transfer reaction mass spectrometry (PTR-MS, Ionicon).

### Animal models

All animal studies were approved by the MIT Institutional Animal Care and Use Committee (protocols 2301000462 and 2203000310) and were conducted in compliance with institutional and national policies. Influenza A A/PR/8/34 (PR8) model was generated by dosing 7-week-old female mice (BALB/c, Taconic) with 25 or 30 μL of Influenza virus through intranasal instillation. Eml4-Alk lung cancer model was generated in 6-week-old female mice (C57BL/6J, Jackson Labs) by intratracheal instillation of 50 μL adenovirus expressing the Ad-EA vector (VQAd Cas9 ALK EML4 072415; Viraquest; 1.5*10^8^ PFU in Opti-MEM with 10 mM CaCl_2_)^35^. These mice are referred to as “Eml4-Alk” mice. The criteria for euthanasia, as dictated by the MIT Committee on Animal Care, was body weight loss of greater than 20% for PR8 mice and 10% for Eml-Alk mice, significant dyspnea, or poor body condition. Animals were monitored daily for PR8 mice and weekly for Eml4-Alk mice throughout studies, and the criteria for euthanasia were not met. Healthy control cohorts were age- and sex-matched mice that did not undergo instillation of influenza virus or adenovirus.

### *In vivo* characterization of vABNs

Peptides for the nanosensors were synthesized by CPC Scientific. The peptides containing an N-terminal cysteine were covalently conjugated to 8-arm 40 kDa PEG maleimide with a tripentaerythritol core (JenKem Technology) at a molecular ratio of 2:1 in water overnight at room temperature. The vABNs were prepared in phosphate buffered saline and delivered to the lungs by intratracheal instillation (50 µL total volume). Breath collection method was reported previously^24^. In brief, at 10- and 30-mins post vABN delivery, mice were placed in 10 mL syringes connected to closed Luer lock stopcock valves for 2 min. Next, 25-gauge needles were connected to the apparatus and inserted through the rubber septum of 12 mL evacuated Exetainers® (Labco). Headspace (10-12 mL) was transferred from the syringe into the Exetainer® and measured by PTR-MS.

### Alectinib treatment of Eml4-Alk mice

Eml4-Alk mice were randomized to receive either control drug vehicle or alectinib (MedChemExpress), at 20 mg/kg prepared directly in drug vehicle, daily by oral gavage for 14 consecutive days. The drug vehicle consists of 10% (v/v) dimethylsulfoxide (DMSO; Sigma Aldrich), 10% (v/v) Cremophor EL (Sigma Aldrich), 15% (v/v) poly(ethylene glycol)-400 (PEG400; Sigma Aldrich), and 15% (w/v) (2-Hydroxypropyl)-β-cyclodextrin (Sigma Aldrich). Mice were monitored daily. Investigators were not blind with respect to treatment.

### Machine learning analysis of multiplexed vABN data

Selection of vABN substrates was performed using analytic pipelines of the Protease Activity Analysis (PAA) package^37^. Analyses of reporter data were performed using Python (v3.9.0) and the scikit-learn^62^ (v1.6) package. For all *in vivo* vABN experiments, z-score standardization was performed on volatile reporter signals in the same mice cohort prior to statistical analysis. Differential vABN signals were subjected to unpaired two-tailed t tests. Multiple comparisons were analyzed by one-way or two-way ANOVAs followed by correction for multiple hypotheses using the Bonferroni method. P_adj_ < 0.05 was considered significant. In Fig. 4, machine learning classification was performed based on the vABN signatures of datasets consisting of paired features (z-scored vABN signals) and labels (the class membership; for example, PR8 or healthy). Classification was performed using a random forest classifier with 100 trees. Estimates of out-of-bag error were used for cross-validation, and trained classifiers were tested on the independent test cohort.

Similarly, in Fig. 5, principal component analysis (PCA) and machine learning classification were performed on datasets consisting of paired features and labels (Eml4-Alk or healthy). Classification of vABN signatures was performed by 25:75 test-train splits of the datasets and trained on gradient boosting classifiers. Hyperparameters for the classifiers were determined by GridSearchCV from the scikit-learn^49,62^ package. A five-fold cross validation was performed on the classifiers, where mean AUC and the F1 score were reported (Supplementary Table 2). Performances of the classifiers were evaluated with the area under the curves (AUCs) of the receiver operating curves (ROC).

## Statistics and reproducibility

PCA and machine learning classifications of vABN data were performed in Python (v.3.9.0) and the scikit-learn^62^ (v1.6) package. All remaining statistical analyses were conducted in Prism 9.0 (GraphPad). Sample sizes, statistical tests, and P-values are specified in figure legends. *In vitro* and *in vivo* experiments were repeated at least twice with similar results.

## Data availability

The data supporting the findings of this study are available within this article and its Supplementary Information. The source data is available at 10.5281/zenodo.15015997.

## Code availability

Code for substrate selections (Figs. 3 and 4) was published as part of the Protease Activity Analysis (PAA) package^29^. The codes for analyzing multiplex data in Figs. 4 and 5 are available at https://github.com/christinewang76/multiplex-mass-analysis.git.

## Acknowledgements

We thank Dr. Ronald Raines (MIT, Cambridge, MA), Dr. Ronald Zuckermann (Berkeley lab, Berkeley, CA), and Dr. William Lubell (Université de Montréal, Montreal, Canada) for valuable feedback and insight on the intramolecular aminolysis and proline chemistry. We thank the Biopolymers & Proteomics and the Preclinical Modeling, Imaging, and Testing cores at the Koch Institute Swanson Biotechnology Center (MIT, Cambridge, MA), with special thanks to Alexander Austin, Heather Amoroso, and Richard Cook for insight on peptide synthesis and Milton Cornwall-Brady for feedback on microCT imaging and analysis. This study was supported, in part, by a Koch Institute Support Grant P30-CA14051 from the National Cancer Institute, a Core Center Grant P30-ES002109 from the National Institute of Environmental Health Sciences, NIH R01 Grant 5R01AI132413-07, the National Institute of General Medical Sciences (T32GM007753 and T32GM144273) (the content is solely the responsibility of the authors and does not necessarily represent the official views of the National Institute of General Medical Sciences or the National Institutes of Health), the Virginia and D.K. Ludwig Fund for Cancer Research, the Koch Institute Frontier Research Program through a gift from Upstage Lung Cancer, the Koch Institute’s Marble Center for Cancer Nanomedicine, and Open Philanthropy. S.T.W and J.D.K. acknowledge fellowship support from the Ludwig Center at MIT’s Koch Institute. C.S.W. acknowledges support from the National Science Foundation Graduate Research Fellowship. S.N.B. is a Howard Hughes Medical Institute Investigator.

