## Supplementary Information for "Engineering Multiplexed Synthetic Breath Biomarkers as Diagnostic Probes"

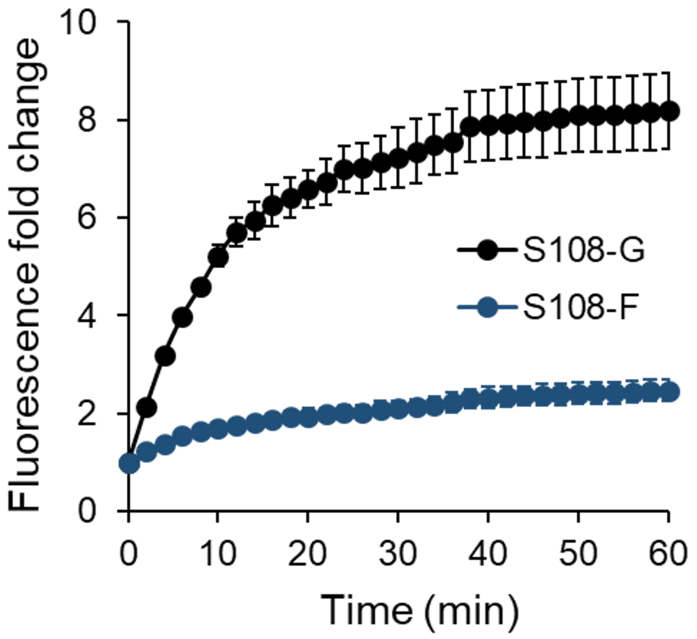


**Fig. S1: Fluorescence kinetics show protease activity is P1' selective.** Two quenched fluorogenic substrates S108-G (FAM-GAANLTRGP-CPQ2) and S108-F (FAM-GAANLTRFP-CPQ2) were designed for sensing trypsin-3 (PRSS3) activities, where P1' reside is either a glycine or a phenylalanine. Both substrates have a canonical cleavage site after P1 arginine. The substrates (5 µM) were mixed with PRSS3 (10 nM) in PBS in a 384-well plate (Corning). Fluorescence kinetics were measured at an excitation and emission of 485 nm and 535 nm, respectively, with a plate reader (Tecan Infinite 200 Pro M Plex) at 37 °C for 1 hr with an interval of 2 min. Fold change was calculated by fluorescence intensity at a given time to that at time = 0 min.

**a**

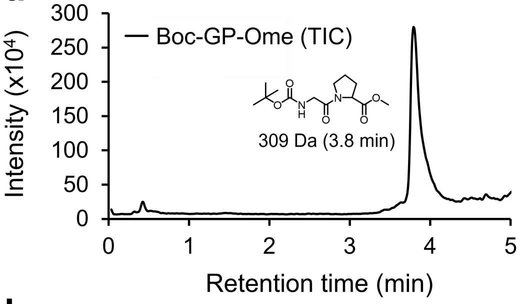


| Extracted mass | Area (x10³) | % |
|---|---|---|
| 309 (286 + Na) | 4798.62105 | 71.6 |
| 187 | 1846.80789 | 27.5 |
| 155 | 60.36321 | 0.9 |

**b**

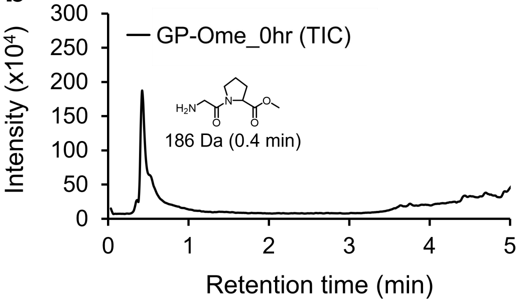


| Extracted mass | Area (x10³) | % |
|---|---|---|
| 309 (286 + Na) | 12.02058 | 0.3 |
| 187 | 4167.61887 | 97.8 |
| 155 | 81.464 | 1.9 |

**c**

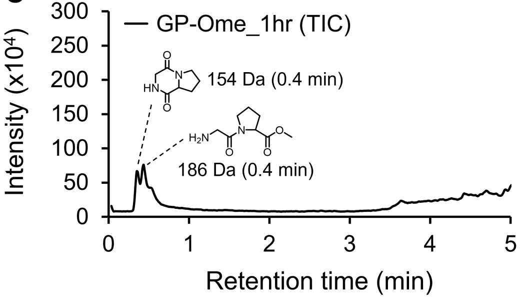


| Extracted mass | Area (x10³) | % |
|---|---|---|
| 309 (286 + Na) | 8.05024 | 2.1 |
| 187 | 86.11381 | 22.4 |
| 155 | 291.04509 | 75.6 |

**Fig. S2: Analysis of aminolysis reaction mediated by chemical cleavage.** Liquid chromatography-mass spectrometry (LC-MS) was used to analyze molecular weights of the reaction products in solution, as described in Fig. 2b. (a) *tert*-butyloxycarbonyl (boc)-GP-Ome (100 ppm), (b) GP-Ome at time = 0 min upon reconstituting in water, and (c) GP-Ome after incubating in water for 1 hr at 37 °C. All samples were prepared in water and adjusted pH to 7. Molecular weight changes show deprotection of boc group by trifluoroacetic acid (b) and conversion to diketopiperazine (c).

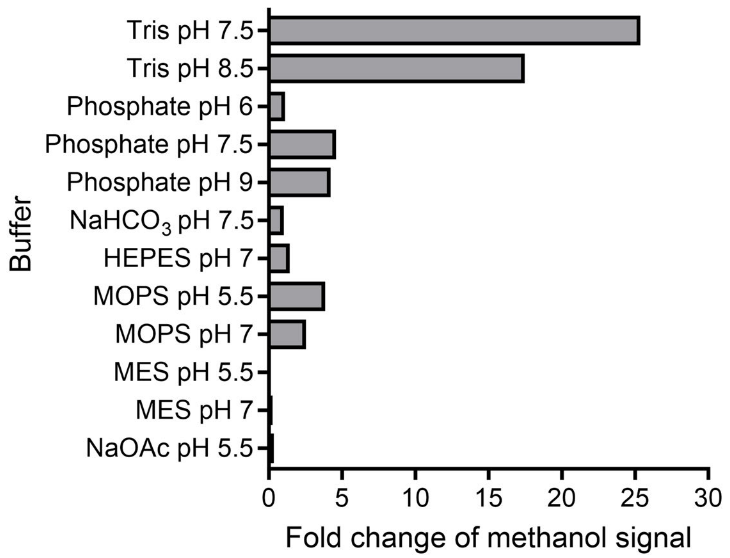


**Fig. S3: Stability of boc-GP-OMe in various buffers.** Boc-GP-Ome was incubated in 50 mM buffers for 1 hr at 37 °C prior to measuring methanol signal in the headspace. These selected buffers are commonly used in activity assays of recombinant proteases. Fold change was calculated by the signal ratios in buffers to water. Release of methanol was observed using tris buffer due to intermolecular aminolysis reaction between tris amine and the ester group of Boc-GP-Ome. Therefore, all tris buffers for protease activity assays were replaced with either HEPES or phosphate buffers.

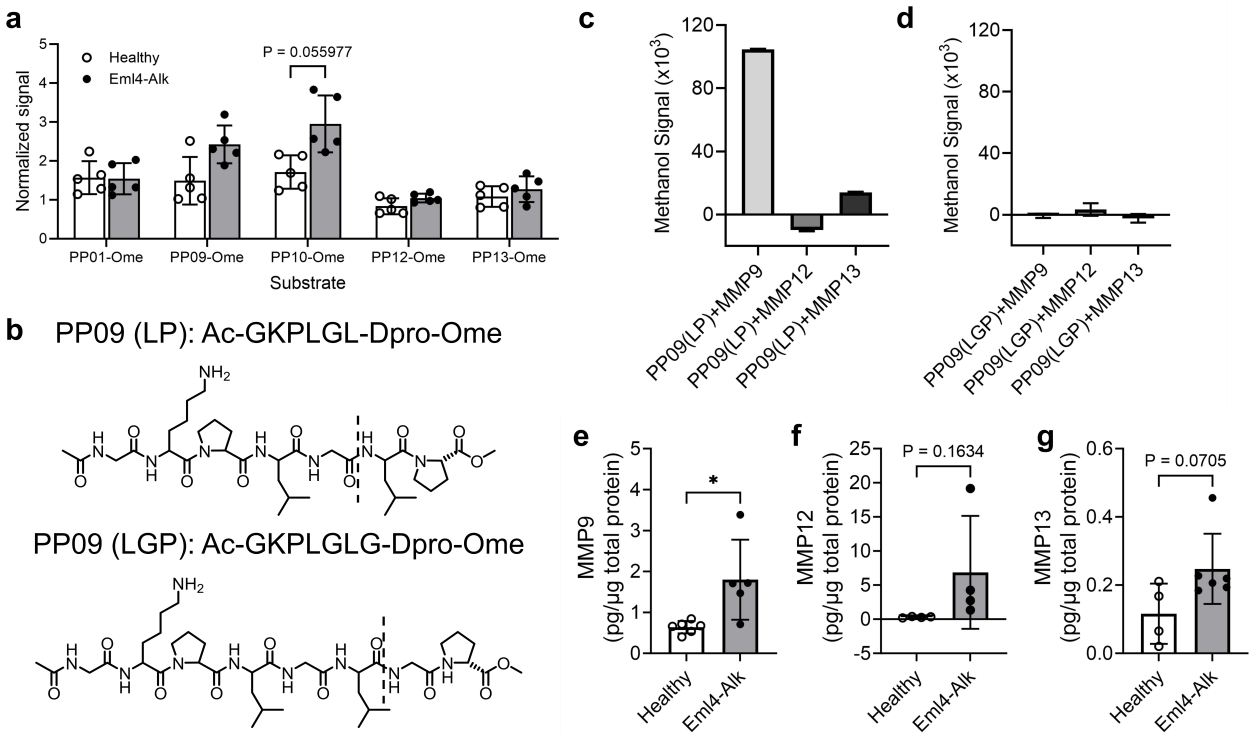


**Fig. S4: *In vitro* assessment of vABN substrates.** (a) Subsets of the urinary probes[1] for Eml4-Alk were re-engineered into vABN substrates. Their ability to differentiate healthy controls and Eml4-Alk mice were assessed using bronchoalveolar lavage fluids (BALF) collected from mice at 7 weeks after tumor initiation. The substrates contain methyl esters that can release methanol following hydrolysis at the canonical sites by target proteases (Table S1). These methanol probes (100 µM) were incubated with BALF (0.1 mg/ml total proteins) overnight at 37 °C. Methanol signals in the headspace were measured by proton-transfer-reaction mass spectrometry (PTR-MS). Statistical significance was calculated by multiple two-sided *t* tests with the Holm-Šídák correction. (b) PP09 contains known MMP-cleavable motifs.[2] To identify its cleavage site by specific types of MMPs, two probes, PP09 (LP) and PP09 (LGP) were generated, which could release methanol following proteolytic cleavage at prescribed sites (dotted lines). (c, d) Methanol signal in the headspace after incubating 100 µM methanol probes with 100 nM recombinant human MMP9, MMP12, and MMP13 for 1 hour at 37 °C. These MMPs were elevated in the transcriptomic analyses of Eml4-Alk lungs[1]. (e-g) ELISAs show MMP levels in BALF collected from healthy controls and Eml4-Alk mice at 7 weeks after tumor initiation.

**Table S1. Sequences of vABNs used in this work.**

1. <u>All substrates for *in vitro* headspace assessments.</u>

| Name | Sequence | Volatile | Target protease | Figure |
|---|---|---|---|---|
| S108-Ome | Ac-GAANLTRGP-Ome | Methanol | Trypsin | 3b, c, d, g, 4c |
| S108(d-pro)-Ome | Ac-GAANLTRGp-Ome | Methanol | Trypsin | 3e, f |
| S108-Od5eth | Ac-GAANLTRGp-Od5eth | Ethanol-$d_5$ | Trypsin | 3h |
| PP01-Ome | Ac-GGPGp-Ome | Methanol | Prolyl peptidase | S4 |
| PP09-Ome | Ac-GKPLGLp-Ome | Methanol | Metalloprotease | S4 |
| PP10-Ome | Ac-GGILSRIp-Ome | Methanol | Serine protease | S4 |
| PP12-Ome | Ac-GPLGMRGp-Ome | Methanol | Serine protease | S4 |
| PP13-Ome | Ac-GGPFGCHAKGp-Ome | Methanol | Serine protease | S4 |

1. <u>All substrates for *in vivo* influenza A in Fig. 4.</u> The sequences of vABNs were obtained from our protease activity analysis database[2].

| vABN | Sequence | Volatile | Target protease |
|---|---|---|---|
| S70 | Ac-CKK(Cy5)-PEG4-GGAIEFD-HFA1 | 2,2,3,3,3-Pentafluoropropylamine | Granzyme B |
| S8 | Ac-CKK(Cy5)-PEG4-GGPLGL-HFA3 | 1H,1H-Perfluoropentylamine | Cathepsin |
| S108 | Ac-CKK(Cy5)-PEG4-GAANLTRGp-Od5eth | Ethanol-$d_5$ | Trypsin |
| S72 | Ac-CKK(Cy5)-PEG4-GGPVPLp-Od7isoprop | 2-propanol-$d_7$ | Metalloprotease |
| S26 | Ac-CKK(Cy5)-PEG4-GRQRRSp-Od3but | 2-butanol-$d_3$ | Furin |

1. <u>All substrates for *in vivo* Eml4-Alk in Fig. 5.</u> The sequences of vABNs were obtained from our previous multiplexed panel for urinary diagnostics of lung cancers[1,3].

| vABN | Sequence | Volatile | Target protease |
|---|---|---|---|
| PP01 | Ac-CKK(Cy5)-PEG4-GGP-HFA1 | 2,2,3,3,3-Pentafluoropropylamine | Prolyl peptidase |
| PP13 | Ac-CKK(Cy5)-PEG4-GGPFGCHAK-HFA3 | 1H,1H-Perfluoropentylamine | Serine protease |
| PP10 | Ac-CKK(Cy5)-PEG4-GPLGMRGp-Od5eth | Ethanol-$d_5$ | Serine protease |
| PP12 | Ac-CKK(Cy5)-PEG4-GGILSRIp-Od7isop | 2-propanol-$d_7$ | Serine protease |
| PP09 | Ac-CKK(Cy5)-PEG4- GKPLGLp-Od3but | 2-butanol-$d_3$ | Metalloprotease |

Note: lower case indicates d-amino acids. 2,2,3,3,3-pentafluoropropylamine (CAS: 422-03-7), 1H,1H-perfluoropentylamine (CAS: 355-27-1), ethanol-$d_5$ (CAS: 1859-08-1), 2-propanol-$d_7$ (CAS: 19214-96-7), 2-butanol-$d_3$ (CAS: 53716-61-3). Ac: acetyl group, Cy5: cyanine 5 fluorophore, and PEG: polyethylene glycol.

**Table S2**. **Cross validation scores for the evaluation of classifiers in Fig. 5.** Five-fold cross validations were performed on the classifiers using Python (v3.9.0) and the scikit-learn[4,5] (v1.6) package.

| Dataset | Mean AUC of folds | F1 score |
|---|---|---|
| 4-week Eml4-Alk (Fig. 5b) | 0.61 ± 0.35 | 0.68 ± 0.22 |
| 5-week Eml4-Alk (Fig. 5b) | 0.83 ± 0.05 | 0.74 ± 0.12 |
| 6-week Eml4-Alk (Fig. 5b) | 0.71 ± 0.3 | 0.72 ± 0.16 |
| 10-week relapse (Fig. 5h) | 0.70 ± 0.19 | 0.76 ± 0.08 |
